# Long-term P fertilization influences microbial carbon use efficiency and soil organic matter decomposition in non-allophanic Andosols

**DOI:** 10.1101/2025.04.29.651165

**Authors:** Wako Koizumi, Timothy J Clough, Soichi Kojima, Tomoyuki Makino, Soh Sugihara, Yoshitaka Uchida, Toru Hamamoto

## Abstract

Phosphorus (P) availability affects soil carbon (C) cycling such as microbial C use efficiency (CUE) and priming effects (PEs). While non-allophanic Andosols are characterized by high organic C content and strong P retention, the effects of different P fertilization regime on C dynamics in these soils remain poorly understood. In this study, we conducted a 20-day incubation experiment using ^13^C-enriched glucose to investigate how different soil P levels (Truog-P: 157 mg P kg^-1^ and 12 mg P kg^-1^) impacted microbial C dynamics in non-allophanic Andosols from contrasting field management practices. Our results showed that soil organic matter (SOM) priming is associated with P fertilization management, with total primed CO_2_-C emissions remaining low in these soils. In the high-P soils, glucose and nitrogen (N) addition resulted in negative PEs, whereas in the low-P soils, the same treatment stimulated microbial SOM mining, resulting in positive PEs. Additionally, higher glucose-derived CUE was found in the high-P soils than in low-P soils after 20 days of incubation. These findings suggest that long-term P fertilization influences both substrate-induced microbial assimilation and SOM decomposition, with P limitation potentially promoting SOM mining which, along with concurrent soil acidity and exchangeable Al toxicity, modulates CUE. This study provides insights for improving C sequestration in non-allophanic Andosols through soil fertility management.

## 1. Introduction

Soil carbon (C) plays a central role in the global C cycle with the potential to mitigate the elevated atmospheric C dioxide (CO_2_) concentration (Jackson et al. 2017). Soil microbes can degrade soil organic C (SOC) emitting it as CO_2_ (Wieder et al. 2013). Increases in the stock of SOC may occur as a result of increased microbial biomass, in response to inputs of C to the soil (Kallenbach et al. 2016; Wang et al. 2021).

Microbial C use efficiency (CUE), the ratio of C allocated to microbial biomass growth to that respired as CO, is positively correlated with SOC levels (Tao et al. 2023). The magnitude of CUE has also been shown to depend on microbial community structure (Maynard et al. 2017), and various abiotic factors, such as soil pH and exchangeable Al, which can alter microbial metabolism and C allocation (Jones et al. 2019; Schroeder et al. 2024), clay content (Islam et al. 2023), nutrient availability (Manzoni et al. 2012; Spohn 2016), and substrate type (Jones et al. 2018). Limited nutrient availability and unbalanced substrate additions may lower CUE by promoting excessive enzyme allocation and respiration, described as overflow metabolism (e.g., Tempest and Neijssel 1992; Russell and Cook 1995; Schimel and Weimtraub 2003; Keiblinger et al. 2010). According to Manzoni et al. (2012), CUE is also influenced by nitrogen (N) and phosphorus (P) availability, since microbes require these nutrients to balance anabolic and catabolic processes. P is an essential nutrient for soil microbes, as it is a key component of DNA, RNA, and has a critical role in phospholipid synthesis, and energy transfer. Depletion of P resources constrains microbial investment in biomass production, shifting the C allocation toward enzyme production, which can lower CUE (Zhai et al. 2022). In addition, limited P availability represents a significant agronomic challenge, as it directly constrains plant productivity and undermines soil health. Microbes compete with plants for P, thereby limiting plant activity by reducing the plant-available P in P-limited soil (Jiang et al. 2024).

An imbalance of available nutrients leads to microbial SOM decomposition and nutrient acquisition, observed as positive priming effects (PEs) (Blagodatskaya and Kuzyakov 2008; Nottingham et al. 2015; Bernard et al. 2022). For example, the addition of a high C:N substrate induces microbial N acquisition from SOM, a process known as ‘N mining’, and positive PEs are found when compared to the addition of low C:N substrates, (Craine et al. 2007; Xu et al. 2024). Immobilization and nutrient mining also applies to P, with high C:P substrates inducing positive PEs to occur under P-limited soil conditions. In some cases, this may also involve the mobilization of P compounds from mineral phases through the mechanisms such as the action of competing ligands, protons, or electron-donating compounds (Spohn 2024; Lieberman et al. 2025). Additionally, P addition to soils may contribute to SOM destabilization through abiotic processes. Inputs of P promotes the desorption of the organic C adsorbed to mineral surface by ligand exchange or the dissolution of the organic matter formed complexes with metal such as aluminum (Al) and iron (Fe) (Kaiser and Zech 1999; Li et al. 2020; Spohn et al. 2022; Mori 2023). Previous studies have indicated that P addition increases the bioavailable C pool and C mineralization, contributing to the SOM decomposition in various soils, including Andosols (Miyazawa et al. 2013; Scott et al. 2015; Spohn and Schleuss 2019; Li et al. 2020).

Considerable research interest has recently focused on P-induced PEs and CUE, and an increasing number of studies examining the effects of P availability on PEs have been conducted using direct P substrate addition, with contrasting results: some showing higher positive PEs but higher CUE (Wang et al. 2025), others with negative PEs (Wang et al. 2014; Sawada et al. 2023) or negative PEs with high CUE (Zeng et al. 2025), or with higher positive PEs but no effects on CUE (Mehnaz et al. 2019). However, short-term additions may not fully capture the sustained microbial and abiotic responses observed under long-term fertilization. Long-term P fertilization in the field provides a more realistic framework than short-term addition for assessing PE responses to exogenous C inputs (Qin et al. 2024). Limited studies using this approach compared to studies using direct P addition have also reported contrasting results. Some observed increased CUE and no significant changes in SOC levels with sustained P input (Li et al. 2025a), while others found enhanced SOM mineralization and positive PEs (Li et al. 2023; Qin et al. 2024). Moreover, Widdig et al. (2020) reported that P fertilization had no effect on CUE. These varied findings emphasize that microbial responses to P availability are highly context-dependent (i.e., microbial nutrient mining hypothesis and microbial stoichiometric decomposition hypothesis, Hessen et al. 2004; Craine et al. 2007). There remains a critical knowledge gap in understanding how P availability influences microbial CUE and PEs across different soil types and the quality or amount of substrate, particularly in soils with unique properties such as Andosols.

Derived from volcanic ash, Andosols are mainly located on the Pacific coast (e.g., Japan, Indonesia, New Zealand, and Chile), but are also present in parts of Europe (e.g., Italy; Iamarino and Terribile 2008; Takahashi 2020; Matus et al. 2024). They are important for soil C sequestration due to their abundance of Al-organic matter complexes which contributes to high SOM stability (Nanzyo 2002; Takahashi and Dahlgren 2016). Andosols are classified according to their significant P retention and are mainly divided into two groups: allophanic and non-allophanic, which are distinguished by the active Al composition (Takahashi and Dahlgren 2016; IUSS Working Group WRB 2022). Non-allophanic Andosols, dominated by Al-organic matter complexes, account for about 30% of Japanese arable Andosols (Saigusa and Matsuyama 1998). Non-allophanic Andosols are characterized by high acidity and high P sorption due to Al-organic matter complexes (Shoji et al. 1987). Thus, non-allophanic variants display greater Al phytotoxicity, which regulates the availability of P, consequently restricting crop growth (Ito et al. 2011). Given the dependence of soil microbial activity on P availability, it is highly expected that microbial CUE is affected by P application. Moreover, long-term P fertilization in the field provides insights into the impacts of sustained P input on C decomposition, which cannot be replicated in short-term laboratory experiments (Qin et al. 2024). However, it remains unknown how CUE responds to long-term P addition in non-allophanic Andosols, and how P-induced SOM changes interact microbial C cycling in non-allophanic Andosols.

In this study, we conducted an incubation experiment to investigate the effect of long-term P fertilization on microbial C utilization in non-allophanic Andosols. We used ^13^C glucose to evaluate the impact of P availability on CUE and soil ^13^C recovery. We hypothesized that: (i) in unfertilized soils, substrate additions would promote P-mining, accelerating the positive PEs; and (ii) higher CUE would be observed in P-fertilized soils, as the nutrient balance would better match microbial requirements compared to unfertilized soils.

## 2. Materials and methods

### 2.1. Soil sampling

Soil samples were collected at 0-10 cm depth in April 2023 from two contrasting fertilization fields at Field Science Center, Graduate School of Agricultural Science, Tohoku University, Japan (38°74’N, 140°75‘E, 190 m above mean sea-level). The soils were classified as non-allophanic Andosols according to the Soil Classification System of Japan (Obara et al. 2011). Two contrasting soils with different P content were sampled (Table 1). The HP soil, which had been P-fertilized since 1987, exhibited a high Truog-P content (157.31 mg P kg¹). The amount of P input from the triple superphosphate and the fused magnesium phosphate varied depending on the cultivation years, ranging from 31 to 361 kg P ha^1^ yr¹. Additionally, cattle manure was applied in certain years. In contrast, the low-P soil (LP) has been cultivated since 2009 without P fertilization, had a significantly lower Truog-P content (12.10 mg P kg¹). Detailed fertilization and cultivation histories are described in Table S1. The HP soil (pH 5.29) had a significantly higher pH than the LP soil (pH 4.49). Correspondingly, the LP soil showed a significantly higher amount of exchangeable Al (21.30 meq kg¹) compared to the HP soil (0 meq kg¹). The collected soils were sieved through a 2 mm sieve, and all visible roots were removed, then soils were stored at 4 until further use.

**Table 1.** Chemical and biological properties in HP and LP soils (mean (SD)). For each property, significant differences between HP and LP soils were determined by independent samples t-tests.

|  | pH<sub>(CaCl2)</sub> |  | Total C |  | Total N |  | Total P |  | C:N | MBC |  | MBP |  | log(MBC:MBP) | Resin P |  | Truog P |  | HCl-P |  |
| --- | --- | --- | --- | --- | --- | --- | --- | --- | --- | --- | --- | --- | --- | --- | --- | --- | --- | --- | --- | --- |
|  |  |  | (g C kg<sup>-1</sup>) |  | (g N kg<sup>-1</sup>) |  | (g P kg<sup>-1</sup>) |  |  | (mg C kg<sup>-1</sup>) |  | (mg P kg<sup>-1</sup>) |  | molar basis | (mg P kg<sup>-1</sup>) |  | (mg P kg<sup>-1</sup>) |  | (g P kg<sup>-1</sup>) |  |
| HP | 5.29 | (0.03) | 88.07 | (3.38) | 5.23 | (0.23) | 2.35 | (0.27) | 16.83 | 1368.71 | (56.55) | 75.06 | (18.31) | 1.68 | 122.39 | (6.31) | 157.31 | (5.07) | 1.41 | (0.029) |
| LP | 4.49 | (0.06) | 91.24 | (1.99) | 5.36 | (0.10) | 1.37 | (0.14) | 17.01 | 1343.65 | (142.09) | 21.31 | (11.63) | 2.31 | 29.13 | (1.33) | 12.10 | (0.31) | 0.54 | (0.063) |
| P values | <0.0001 |  | n.s |  | n.s |  | <0.01 |  | <0.05 | n.s |  | <0.01 |  | <0.05 | <0.001 |  | <0.0001 |  | <0.0001 |  |

MBC, Microbial biomass C; MBP, microbial biomass P; MBC: MBP, their molar-based ratio; Resin P, Resin-extractable P. Truog P and HCl-P
represent Truog-extractable P and 1M HCl-extractable P, respectively.

### 2.2. Experimental design

Prior to the experiment, the sieved soils were pre-incubated in the dark at 25L for 14 days and maintained at a gravimetric moisture content of 45% (w/w). Pre-incubated soil was placed in plastic bags (each equivalent to 100 g dry weight). Soil samples received a substrate solution containing either: (1) water (“Control”), (2) ^13^C-enriched glucose (“Glc”), or (3) ^13^C-enriched glucose and (NH_4_)_2_SO_4_ (“Glc+N”). There were 24 samples in total (two soils × three glucose levels × four replicates). The rate of added ^13^C-enriched glucose (4 atom%, D-GLUCOSE (U−^13^C_6_, 99%), Cambridge, 450 mg C kg^-1^ soil) was set to be 25 to 50% of the MBC in order to increase microbial activity while avoiding exponential microbial growth and shifting microbial community composition (Blagodatskaya and Kuzyakov 2008; Sawada et al. 2023). The concentration of added mineral N (61 mg N kg^−1^ soil) was determined according to mean microbial C:N:P ratios, 60:7:1 (Cleveland and Liptzin 2007). The experiment was then started by wetting to 55% gravimetric soil moisture content with each substrate solution.

### 2.3. CO_2_ emission

For measurement of soil respiration, each treatment was placed in a 450 mL airtight glass jar with a CO_2_ alkali trap (5 mL of 1 M NaOH) and incubated at 25L for 20 days. Ten mL of 0.01 M HCl solution was also placed in the glass jar to maintain the soil moisture. The collected NaOH solutions received 8 mL of 1 M SrCl_2_ to form SrCO_3_ precipitate, and the resulting mixture of precipitate and solution was titrated using a 1 M HCl solution (Zibilske 1994; Qiao et al. 2014). The collected NaOH solutions received 8 mL of 1 M SrCl_2_ to form SrCO_3_ precipitate and this was titrated using a 1 M HCl solution. After titrating, the precipitates were washed three times with 30mL water and then dried at 105L. The dried SrCO_3_ was packed in tin capsules to analyze for ^13^C enrichment on DELTA V plus isotope ratio mass spectrometer (IRMS, Thermo Fisher Scientific, Bremen, Germany). The calculation of glucose-derived CO_2_-C_glc_ was adapted from Mehnaz et al. (2019) and Brant et al. (2006):

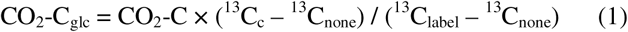

Where CO_2_-C is the cumulative CO_2_-C respired from the soil with glucose addition; ^13^C_c_, ^13^C_none_ and ^13^C labels are the ^13^C atom% values of the CO_2_-C measured in the NaOH solutions with and without glucose addition and the added glucose substrate, respectively. Soil-derived CO_2_-C (CO_2_-C_soil_) in glc and glcN treatments was calculated as the difference between the total (CO_2_-C) and glucose-derived CO_2_-C_glc_.

### 2.4. Soil microbial biomass C and P

Soil microbial biomass C (MBC) was determined with a modified chloroform fumigated method (Beck et al. 1997). Briefly, two samples of 5 g dry soil of each sample with and without 24 h fumigation using chloroform were extracted with 25 mL of 0.05 M K_2_SO_4_ and analyzed for total organic C using a TOC-TN analyzer (TOC-V CPH/CPN, Shimadzu, Japan). The MBC was calculated from the difference in total organic C concentrations in samples with and without fumigation divided by an extraction efficiency of 0.45 (Beck et al. 1997). The ^13^C atom% in each extract was measured on an IRMS (Thermo Fisher Scientific, USA). The ^13^C atom% of the microbial biomass (^13^C_mic_) and the glucose-derived C in the microbial biomass (MBC_glucose_) were also calculated by the following equation (Mehnaz et al. 2019):

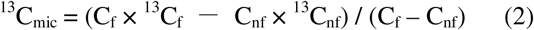

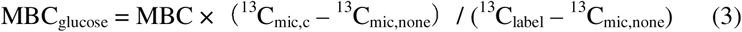

where C_f_ and ^13^C_f_ are the amount of C and ^13^C atom% in the fumigated extracts, and C_nf_ and ^13^C_nf_ are those in the non fumigated, respectively.

Organic C within K_2_SO_4_ extracts from non-fumigated soil was denoted as Dissolved organic C (DOC). The terms ^13^C_mic,c_ and ^13^C_mic,none_ are the ^13^C_mic_ values with and without glc addition.

Soil microbial biomass P (MBP) was measured using an anion exchange membrane method according to Kouno et al. (1995). Briefly, soil samples (1 g dry mass equivalent) were shaken and extracted in a 20 ml mixture of deionized water and resin for 16 h. For fumigated samples only, 2 ml of hexanol was added instead of chloroform used in the method of Kouno et al. (1995). The resin was then removed and the phosphate was recovered by shaking for 1 h in 20 ml of 0.5 M HCl. We added 25 and 40 mg P kg^−1^ depending on P contents as spike considering the P adsorption in LP and HP soils, respectively (Brookes et al. 1982). The extracted P was determined with the malachite green colorimetric method (Ohno and Zibilske 1991; Jeannotte et al. 2004; Song et al. 2019). The extracts were diluted appropriately to avoid interference with HCl (D’Angelo et al. 2001) and analyzed by absorbance measurement at 630 nm using a microplate reader (iMark, Bio-Rad Laboratories, USA). The MBP was calculated from the difference in extracted P concentrations between samples with and without fumigation divided by 0.40 (Brookes et al. 1982).

### 2.5. Soil P measurements

Phosphate within non-applied hexanol extracts for analysis of MBP was denoted as Resin extractable P (Resin P). Resin P represent a part of labile P pools (Erinle et al. 2020) and avoid fixation of P by soil colloid during the extraction in Andosols (Kouno et al. 1995). However, we note that accurately assessing P availability in Andosols with high P sorption capacity using a single chemical extraction method is difficult (Fixen and Grove 1990; Sugito and Shinano 2013). 1M HCl extractable P described by DeLuca et al. (2015) was also measured. It primarily represents P mobilized by protons released by microbes or plants mainly from apatite and other recalcitrant inorganic P forms, and also partially extracts Al/Fe-bound P and phosphate which is weakly adsorbed to clay particles or present in inorganic precipitates (Kuo 1995; Pingree et al. 2017; Wuenscher et al. 2015; Li et al. 2019). This fraction was denoted as HCl-P. It was measured by extracting 0.5 g of dry mass equivalent soil with 10 ml of 1 M HCl for 3 h at 200 rev min^−1^ at room temperature. After filtering, the inorganic P was determined by the malachite green method at 630 nm using a microplate reader (iMark, Bio-Rad Laboratories, USA) as for the MBP analysis.

### 2.6. Inorganic N measurements

Inorganic N (IN) was determined as the sum of NHLL-N and NOLL-N. Soil samples (5 g dry mass equivalent) from the 2- and 20-day incubation periods were extracted with 25 ml of 1 M KCl by shaking for 30 minutes. The concentrations of NHLL-N and NOLL-N in the filtrates were analyzed using a flow injection analyzer (AQLA-700, Aqualab Co., Ltd., Japan), with NHLL-N measured by absorbance at 630 nm and NOLL-N by absorbance at 540 nm as described by Hamamoto and Uchida (2015).

### 2.7. Substrate-derived C use efficiency and Priming effects

The calculations of glucose-derived C use efficiency (CUE) and priming effects (PEs) were adapted from Mehnaz et al. (2019). While the CUE measured in this study strictly represents glucose use efficiency, we used term “glucose-derived CUE” to maintain consistency with similar previous studies (e.g., Cui et al. 2022; Karhu et al. 2022, Kalu et al. 2024, Xiao et al. 2021). Microbial glucose-derived CUE (CUE_glucose_) of the added glucose was calculated using the following equation:

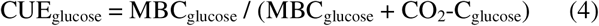

The PEs were calculated as the increase in native soil organic C released after substrate application also using the following equation (Mehnaz et al. 2019):

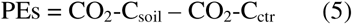

Where CO_2_-C_soil_ and CO_2_-C_ctr_ are the soil-derived CO_2_-C respired from the soil with glucose addition and the cumulative CO_2_-C respired from the soil without glucose addition, respectively.

### 2.8. Soil extracellular enzyme activities

The potential activity of acid phosphomonoesterase (EC 3.1.3.2), P-acquiring extracellular enzyme activity (EEA), was measured fluorometrically with methylumbelliferone (Bell et al. 2013). Briefly, soil samples (1.5 g dry mass equivalent) with 45 mL of deionized water were shaken for 30 minutes (Deng et al. 2011; Li et al. 2025b). 800 μL of mixed soil slurry was taken into a deep-well plate, then 200 μL of 1 mM substrate (4-Methylumbelliferyl phosphate) was added and incubated at 35 °C for 1.5 hours. After incubation samples were centrifuged for 3 minutes at 2500 xg and moved to the 96-well black microplate. Fluorescence intensity was measured using a Varioskan ALF multi-mode microplate reader (Thermo Scientific, USA) at 345 nm excitation and 450 nm emission. EEA was expressed as nmol g^−1^ h^−1^.

### 2.9. Statistics

All statistical analyses were performed with R (version 4.2.2; R Core Team. 2023). We conducted a two-way analysis of variance (ANOVA) including the interaction between soils and substrates to assess their effects on the total CO_2_-C, glucose-derived and soil-derived CO_2_-C using linear model (Bates et al. 2015). We also conducted three-way ANOVA including the interaction between soils and substrates, soils and time, and substrates and time to assess their effects total MBC, glucose-derived and soil-derived MBC, DOC, MBP, HCl-P, Resin P, and EEA using linear model. Prior to the analysis, the datasets were rank-transformed to meet the assumptions of ANOVA. Model was validated by examining residuals using the DHARMa package (Hartig 2022). The data were further analyzed using Tukey’s HSD test to find pairwise differences between soil type and substrate combinations at α = 0.05 using multcomp package (Hothorn et al. 2008). The effects of soil P fertilization on each result, N addition on PEs was determined by independent samples t-tests.

## 3. Results

### 3.1. CO_2_ emission

The HP soil had a higher cumulative CO_2_ emission (339.3 mg C kg^-1^ soil) than the LP soil (308.6 mg C kg^-1^ soil) and there were significant differences when averaged across substrate treatments during the incubation period (Fig. 1 and S1). Substrate additions significantly stimulated the CO_2_ emissions from both soils at day 1 immediately after the incubation (Fig. S1 and Table S3). This initial increase was primally due to the glucose-derived CO_2_ emission, which peaked at 161.2 mg C kg^-1^ in the HP soil and 154.5 mg C kg^-1^ in the LP soil across substrate treatments (Fig. S1). Among substarte treatments, the glc treatments had significantly higher cumulative CO_2_ emissions than the glcN treatments in each soil (Fig. 1 and Table S3).

**Fig. 1.**
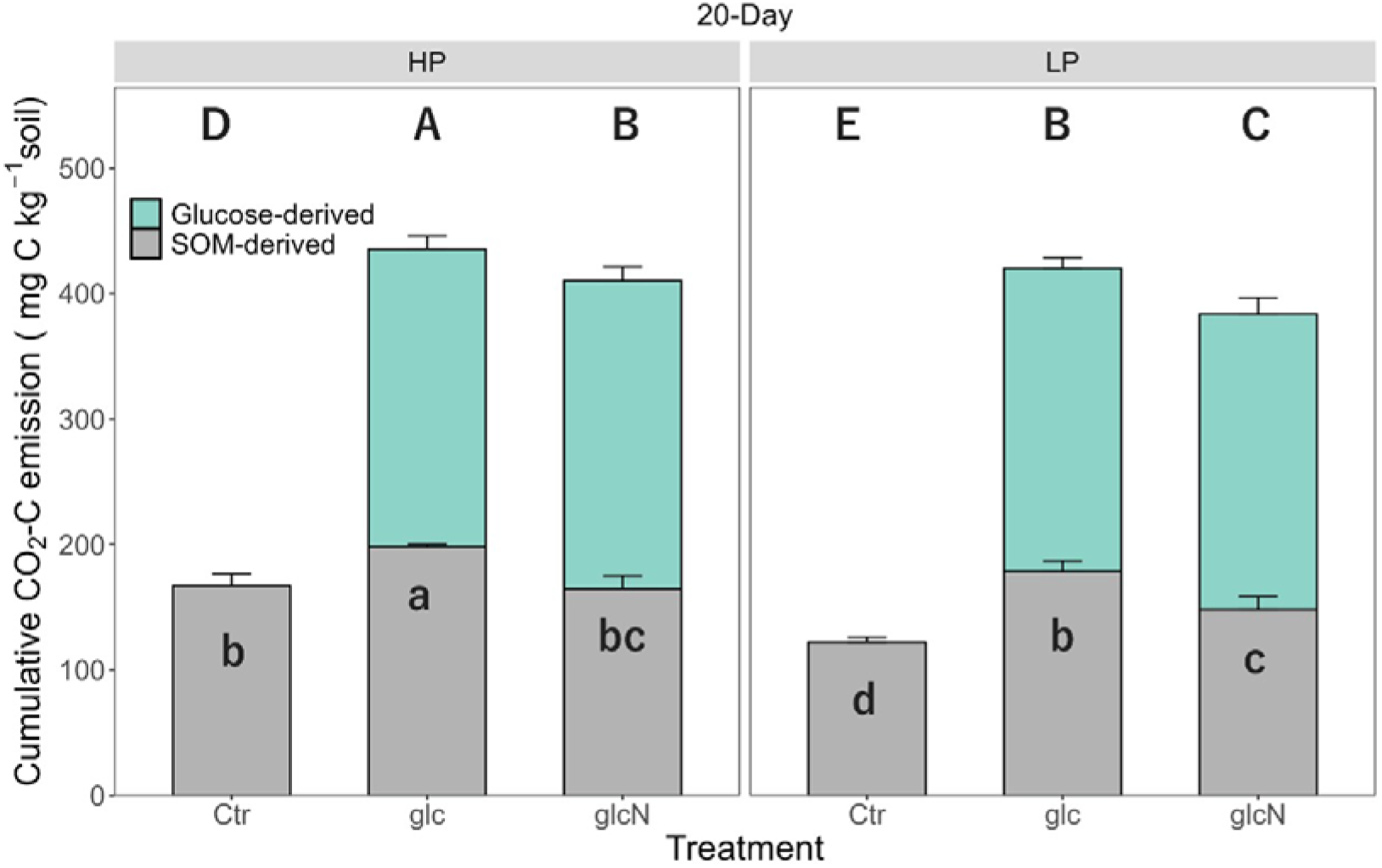
Cumulative CO_2_ emissions (mg C kg^-1^ soil) over 20-days incubation. Treatments include soil without any amendment (Ctr), soil with ^13^C-enriched glucose (glc), and soil with ^13^C-enriched glucose and N substrates (glcN). Different color bars show cumulative CO_2_ emissions (mean ± SD, n=4, but n=3 in LP Ctr due to an operational error during the experiment) derived from added glucose (light green) and soil organic matter (grey). Uppercase and lowercase letters indicate significant differences (p < 0.05) in total and SOM-derived CO_2_-C among treatments, respectively.

The cumulative SOM-derived CO_2_ emission was higher in the HP soil than the LP soil averaging across the substrate treatments (p<0.05, Fig. 1). Substrate additions increased SOM-derived CO_2_ emission in each soil type 6 days after incubation (p<0.001, Table S3). In the HP soil, the glc treatment increased the SOM-derived CO_2_ emissions, while in the LP soil, glc and glcN treatments significantly increased the SOM-derived CO_2_ emissions.

In contrast to the total and SOM-derived CO_2_ emissions, the cumulative glucose-derived CO_2_ emission showed no significant differences among soils and substrate treatments throughout the incubation period (Fig. S2).

The PEs during incubation were lower in the HP soil than the LP across the treatments (p<0.05 at 1, 6, 8 days of incubation, p<0.01 and p<0.001 at 2 days and 12, 20 days of incubation, respectively) (Fig. 2, Tables S3 and S4). The glc addition had significantly higher PEs compared to the glcN addition in both soils. The addition of glc and glcN in the LP soil showed positive PEs. However, the effect of substrate treatments on PEs in the HP soil contrasted with the LP soil; the glc addition showed a positive PEs, but glcN addition showed negative PEs. The total primed CO_2_-C as a % of added glucose C ranged from –0.71 to 12.88 % (Table S4).

**Fig. 2.**
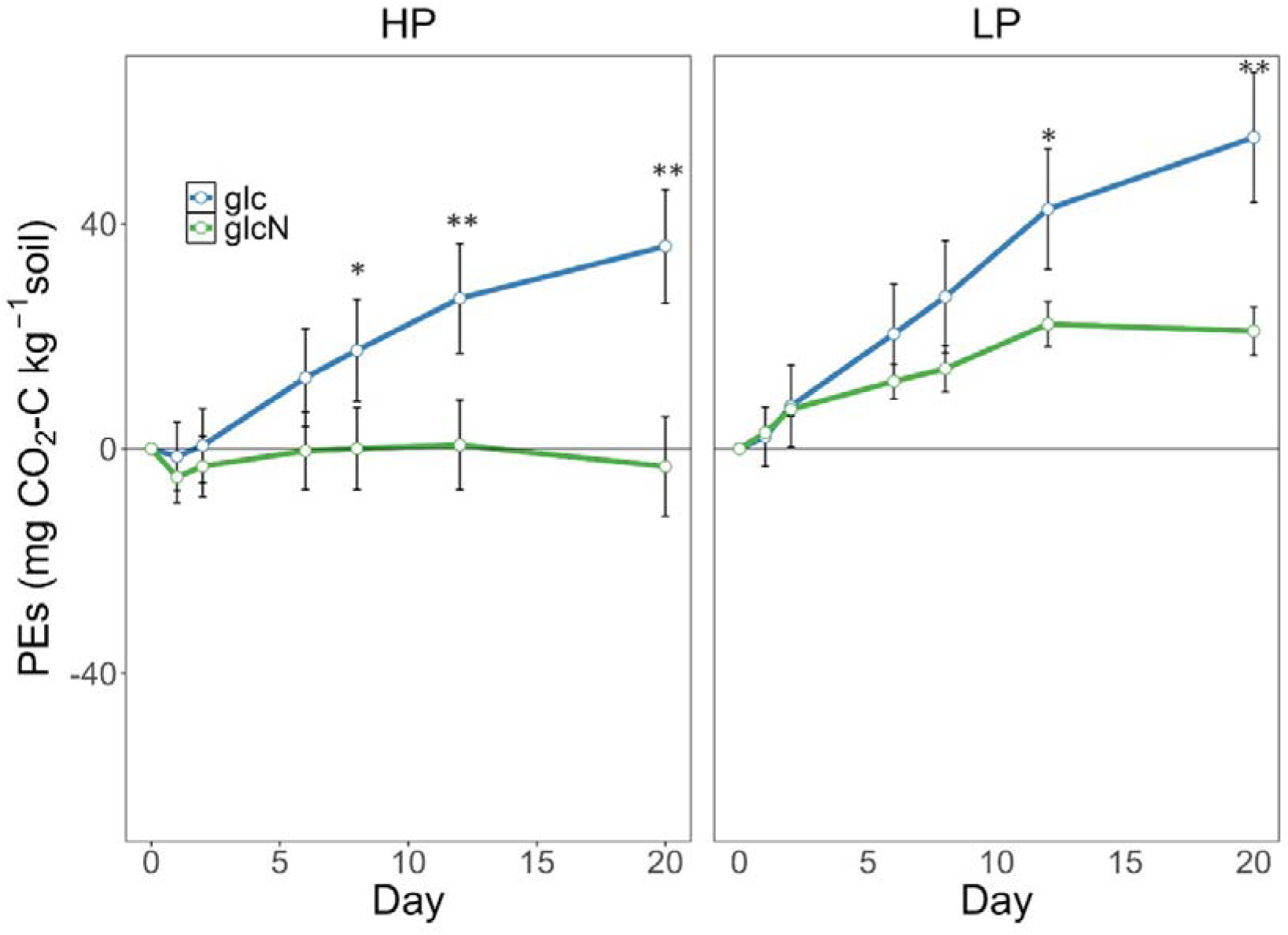
The priming effects (PEs) in HP and LP soils over 20 days. Treatments include soil with ^13^C-enriched glucose (glc) and soil with ^13^C-enriched glucose and N substrates (glcN). Green and blue lines show the priming effect (mean ± SD, n = 4) in glc and glcN, respectively, for each soil. * (p <0.05), ** (p <0.01) and blank (not significant) indicate significance level of N addition effect for each soil separately.

### 3.2. Microbial biomass, soil chemical properties, and soil enzyme activity

The HP soil had significantly higher total MBC (MBC_Total_) than the LP soil at 20 days of incubation, whereas no significant differences were observed after 2 days of incubation (Fig. 3, Table S5 and S6). Substrate additions did not affect MBC_Total_ in either soil at either 2 or 20 days of incubation (Table S5 and S6). For the SOM-derived microbial biomass C (MBC_SOM_), substrate additions significantly decreased MBC_SOM_ in the HP soil compared to the control at 2 days of incubation (Fig. 3, Table S5 and S6). In contrast, the LP soil showed no differences in MBC_SOM_ among substrate treatments at 2 days of incubation. Glucose-derived microbial biomass C (MBC_Glc_) showed no significant differences among soils and substrate treatments at 2 days of incubation (Fig. 3, Table S5 and S6). However, at 20 days of incubation, MBC_Glc_ was higher in the HP soil than in the LP soil. In both soils, MBC_Glc_ was significantly lower at 20 days of incubation compared to 2 days. The P-acquiring enzyme activity was higher in HP soils than in LP soils (Table S5 and S6). The addition of GlcN significantly increased the P acquire enzyme activity in LP soils at 2 and 20 days of incubation.

**Fig. 3.**
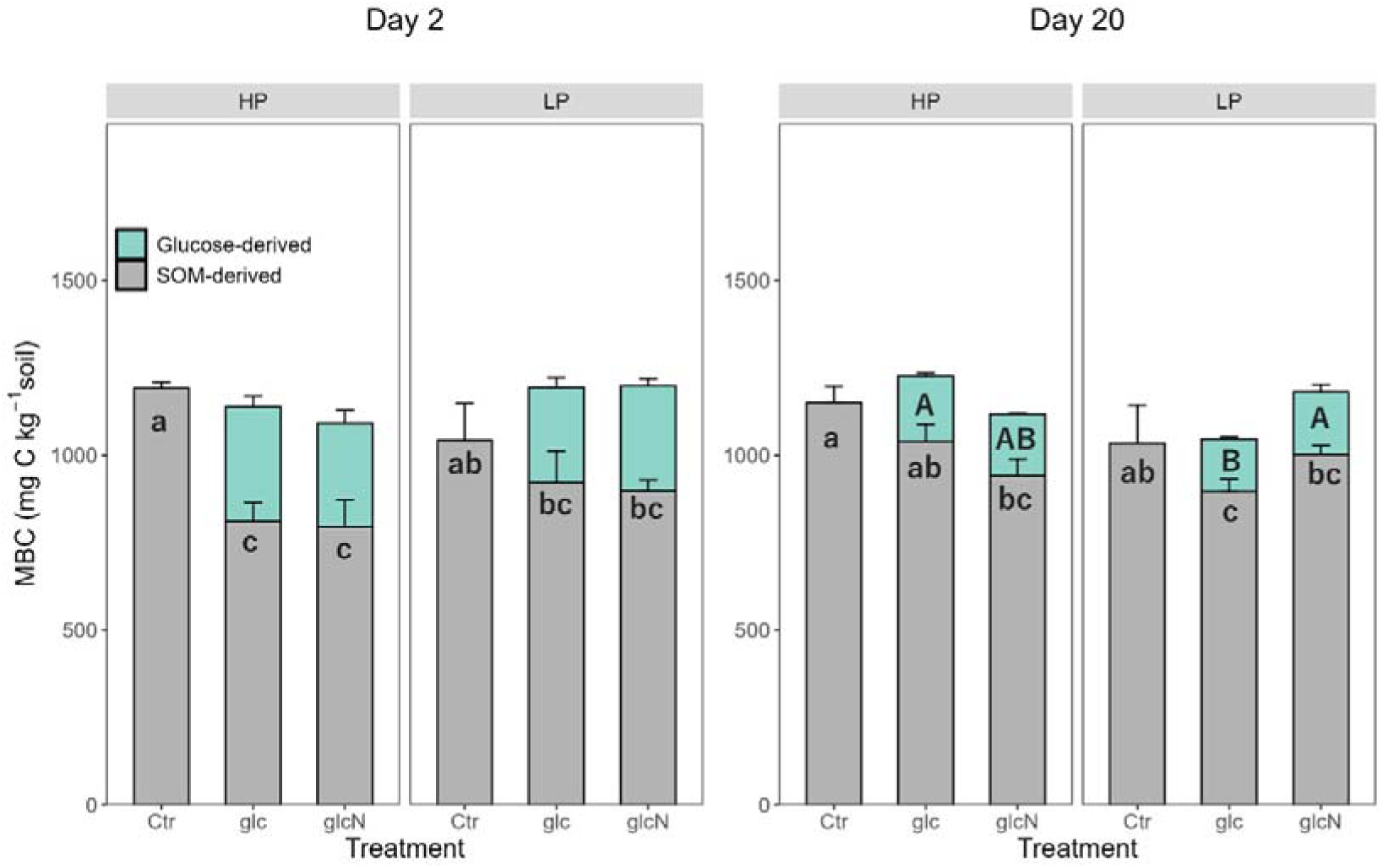
Microbial biomass C (MBC) (mg C kg^-1^ soil) in 2 and 20-days incubation. Treatments include soil without any amendment (Control), soil with ^13^C-enriched glucose (glc), and soil with ^13^C-enriched glucose and N substrates (glcN). Different color bars show cumulative MBC (mean ± SD, n =4, but n =3 in LP Ctr due to an operational error during the experiment) derived from added glucose (light green) and soil organic matter (grey). Uppercase and lowercase letters indicate significant differences (p <0.05) in glucose-derived MBC and SOM-derived MBC among substrate treatments and soils at each sampling time, respectively.

The HP soil had significantly higher MBP and a lower log ratio of MBC:MBP compared to the LP soil at 2 and 20 days of incubation (Table S5 and S6). However, substrate additions had no significant effect on MBP in the both soils during the incubation period.

Soil pH, Resin P, and HCl-P were consistently higher in the HP soils than in the LP soil during the incubation period (Table S6 and S7). Resin P increased from 0 to 2 days of incubation in HP soil (p <0.001), while there was no significant difference in LP soil (p =0.065). Then Resin P decreased from 2 to 20 days in LP soil (p <0.001), whereas no significant difference was found in HP soil (p =0.41).

### 3.3. Microbial C use efficiency

Glucose-derived microbial C use efficiency (CUE) showed no significant differences between the two soils or substrate additions at 2 days after incubation (Table 2). However, at 20 days after incubation, glucose-derived CUE in the HP soil was higher than in the LP soil (p <0.05).

**Table 2.** C use efficiency (CUE) in the HP and LP soils at 2 and 20 days after incubation (mean (SD)). Different letters indicate significant References (p <0.05) among treatments for each soil type.

| CUE |  |  |  |  |  |  |
| --- | --- | --- | --- | --- | --- | --- |
| Soil | Substrate | 2-day |  | 20-day |  |  |
| HP | glc | 0.65 | (0.025) | 0.44 | (0.015) | a |
|  | glcN | 0.61 | (0.038) | 0.42 | (0.009) | ab |
| LP | glc | 0.62 | (0.035) | 0.39 | (0.012) | b |
|  | glcN | 0.63 | (0.012) | 0.43 | (0.025) | a |
| **ANOVA P values** |  |  |  |  |  |  |
| Soil |  | n.s |  | <0.05 |  |  |
| Substrate |  | n.s |  | n.s |  |  |
| Soil × Sub | | n.s | | <0.01 | | |

The HP soil with glc showed the highest glucose use efficiency, while the LP soil with glc had the lowest glucose use efficiency at 20 days after incubation.

## 4. Discussion

Our results show that contrasting long-term P fertilization affected on the microbial respirations. The high-P soils had a significantly higher cumulative CO_2_ emissions in the control compared to the low-P soils, and this trend also remained consistent regardless of the glc or glcN addition. This finding suggests that P input potentially enhanced the microbial activities due to increasing available P and could also promote the desorption of the organic C, thereby increasing available C pools (Kaiser and Zech 1999; Miyazawa et al. 2013; Scott et al. 2015, Spohn and Schleuss 2019, Li et al. 2020; Spohn et al. 2022).

PEs were also influenced by P fertilization (Fig. 2). Notably, N-mining was suggested in both soils by the higher soil-derived COL emissions with glucose-only addition compared to glucose+N addition (Li et al. 2017). However, the overall response of PEs to substrate addition clearly contrasted with soil P status. Specifically, when N was not limited (i.e., in the glucose+N treatment), negative PEs were found in the high-P soil, whereas positive PEs were observed in the low-P soil. Combined with the significantly increased phosphatase activity in the low-P soil (Table S5), this result indicates that the positive PEs are associated with SOM degradation driven by P-mining, supporting our first hypothesis. In addition, the low-P soils had consistently lower Resin P and higher MBC:P ratios compared to the high-P soils (Table 1, S6, and S7). Under nutrient-imbalanced conditions, microbes invest abundant elements into decomposing SOM to obtain limited nutrients, thereby maximizing their productivity (Allison and Vitousek 2005; Mooshammer et al. 2014). Moreover, MBP levels in the low-P soil remained stable throughout the incubation period, possibly because positive priming induced P-mining and maintained microbial P (Spohn 2016).

In agreement with our second hypothesis, the P fertilization was also associated with CUE. After 20 days of incubation, the high-P soils showed significantly higher glucose-derived CUE than the low-P soils (Figs. 1, 3 and Table 2). These long-term CUE measurements (>48h) suggest the efficiency of C stabilization through the recycling of the microbial necromass and exudates (Geyer et al. 2016; Kalu et al. 2024), whereas the short-term incubation CUE measurements (<48 h) may reflect the substrate utilization for gross production (Brant et al. 2006; Geyer et al. 2016). After 20 days of incubation, in the low-P soil, microbes allocated more energy to catabolic pathways to acquire P, which in turn reduced C available for growth and resulted lower glucose-derived CUE than in the high-P soil (Bernard et al. 2022). These findings suggest that differences in P fertilization between the two soils are a key factor explaining the variation in CUE and PEs.

The two soils used in this study significantly differed in pH and exchangeable acidity due to long-term P fertilization, both of which are important factors influencing microbial activity. The lower pH throughout the incubation period and higher exchangeable Al in the low-P soils compared to the high-P soils likely suppressed the microbial respirations and activities (Table1 and S7). In general, higher PEs occur in soils with neutral pH compared to the soils with acidic pH because the microbial activity, the community structure, and the enzyme synthesis are more pronounced (Blagodatskaya and kuzyakov 2008; Blagodatskaya and Anderson 1998). However, previous study suggested that greater positive PEs observed in strongly acidic soils are associated with the size of labile C pools and net increase in MBC (Aye et al. 2018). In this study, although MBC showed no significant differences among soils, higher DOC concentrations were shown in the low-P soils across the treatments, which corresponded with the greater primed CO_2_ emissions observed (Fig. 3 and Table S5). These findings indicated a potential role of low pH in affecting DOC pools and microbial resource accessibility. In contrast, in the low-P soils, MBP showed significant lower than the high-P soils, and as discussed above, the higher MBC:P ratio likely reflects SOM degradation induced by P-mining.

In addition to PEs, soil pH is also important for understanding the CUE results; however, after 2 days of incubation, glucose-derived CUE showed no significant differences among soils (Table 2). It is possible that the effect of P fertilization on microbial assimilation of substrates (i.e., short-term CUE) after 2 days of incubation was offset by: i) availability of organic C; ii) the soil pH, both specific to non-allopnanic Andosols. Soil C content is one of the regulating factors of microbial assimilation (Kamble and Bååth 2014); Andosols have a large amount of organic C content due to Al-soil organic matter complexes (Takahashi and Dahlgren 2016). Moreover, it has been suggested that non-allophanic Andosols have higher organic C contents than other Andosols (Nanzyo et al. 1993). Previous studies showed that microbial growth and CUE are strongly related to C availability rather than N and P availability (Wang et al. 2021; Morris et al. 2021; Jiang et al. 2025). In this study, we found that the high-P and the low-P soils contain approximately 10% TC, this high C content may have influenced glucose-derived CUE after 2 days of incubation. Moreover, soil pH can alter the composition of microbial communities, and consequently affect CUE (Manzoni et al. 2012; Zhen et al. 2019). Rousk et al. (2011) reported that fungal growth is suppressed at pH_(H2O)_ >5.5, contrasted with bacterial growth limited at pH_(H2O)_ <5.5. Keiblinger et al. (2010) reported that fungi induced higher CUE than bacteria, and fungal CUE was affected by C availability and bacterial CUE limited by N and P availability, respectively. Thus, it is possible that the microbial community composition was adapted to low pH conditions and the high C content of the soil and these factors may have contributed to the marginal effect of P fertilization on glucose-derived CUE in the non-allophanic Andosol.

Jones et al. (2019) reported that low pH and associated exchangeable Al decrease CUE through metabolic shifts and greater C diversion into maintenance and catabolic processes, rather than through a reduction in the ability to utilize labile C. In our study, the amount of exchangeable Al was 0 meq kg^-1^ in the high-P soil and 21.30 meq kg^-1^ in the low-P soil, respectively. It is possible that the lower glucose-derived CUE in the low-P soils after 20 days of incubation, when applied C substrate was largely consumed, was due to the toxicity of exchangeable Al, as reflected in the differing amounts of Al^3+^, in addition to low soil P level. Although Jones et al. (2019) focused on short-term effects (2-72 h), our study suggested that these factors possibly affect CUE over longer time scales than several hours. The absence of a difference in CUE between the high-P and the low-P soils on 2 days of incubation, however, may have been due to the influence of organic C availability, as mentioned in the previous paragraph. Therefore, the low CUE observed in the low-P soils is likely attributable to C investment for P acquisition and the response to acid and Al stresses.

Our results showed lower primed CO_2_-C of added glucose C in non-allophanic Andosols compared to previous studies. In the current, the values ranged from –1 to 13 % in the non-allophanic Andosols over 20 days incubation compared to values of 28 to 420 % for other soil types (Table S4) (Shahbaz et al. 2018; Mehnaz et al. 2019; Sawada et al. 2023). This comparison is valid as the values were calculated from data between 18 to 42 days of the entire incubation period. In contrast, the size of the MBC pool was comparable or even larger than the size in these studies (100-1190 mg C kg^-1^ soil), suggesting a higher total microbial biomass including potentially active microbes. This lower PEs in the non-allophanic Andosols were possibly due to the abundance of Al-organic matter complexes. The stability of SOM depends on Al-organic matter complexes and soil acidity (Miyazawa et al. 2013). These complexes promote the soil aggregates formation and are also suggested to physically suppress the microbial SOM decomposition (Matsui et al. 2016). Thus, it is suggested that the low PEs reflect SOM stability in non-allophanic Andosols.

Our findings indicate that maintaining C stocks in non-allophanic Andosols depends not only on adequate long-term P fertilization but also on appropriate regulation of soil acidity. However, we acknowledge that our experiment did not directly represent the temporal Al toxicity effects on CUE, and we could not statistically separate the individual contributions of these overlapping factors to the observed responses. Future studies need to demonstrate these causal relationships by employing controlled manipulations of soil pH and investigate the relationships between Al bioavailability and C dynamics at different soil P levels.

## 5. Conclusion

This study demonstrated that substrate-induced SOM decomposition in non-allophanic Andosols is strongly influenced by the long-term P fertilization. The addition of glucose with N substrate promoted positive PEs under low P conditions, while negative PEs occurred under high P conditions. Throughout the incubation, PEs in the low-P soils were greater than in the high-P soils. Additionally, in the low-P soils, substrate additions did not affect MBP and the ratio of MBC:P throughout the incubation period. These results suggest that a lack of P affected the impact on SOM decomposition due to the maintaining microbial P. Greater CUE was observed in the high-P soils than in the low-P soils after 20 days of incubation, with no difference after 2 days of incubation. These results indicate that under P-limited conditions combined with soil acidity and exchangeable Al, the microbial allocation of C for P acquisition regulates substrate-induced microbial assimilation and SOM decomposition. Future research needs to elucidate the interactive effects of P availability, soil pH, and exchangeable Al on microbial C dynamics in non-allophanic Andosols.

## Author’s Contributions

W.K., T.M., and T.H. planned and designed the research. W.K. and T.H. conducted the incubation experiments. W.K. and T.H. analyzed the data, wrote the main manuscript, and prepared the figures and tables. All authors reviewed the manuscript.

## Acknowledgments

This work was financially supported by the JSPS Grant-in-Aid for JSPS Fellows (23K14056), and the Yamaguchi Educational and Scholarship Foundation. This study was supported by Support system for young researchers to use research equipment in Tohoku University.

