## Supplemental figures for "Long-term P fertilization influences microbial carbon use efficiency and soil organic matter decomposition in non-allophanic Andosols"

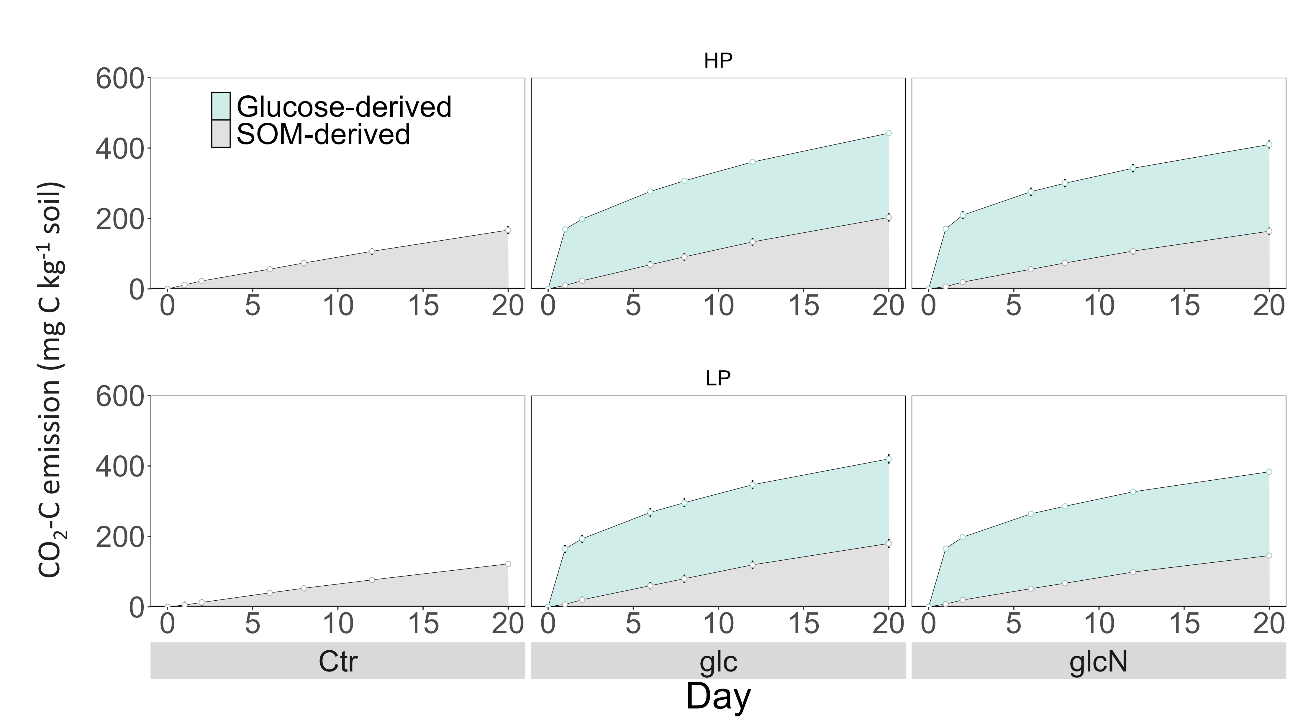

Fig. S1. Cumulative CO_2_ emissions (mg C kg^–1^ soil) from HP and LP soils over 20 days incubation. Treatments include soil without any amendment (Control), soil with ^13^C-enriched glucose (glc), and soil with ^13^C-enriched glucose and nitrogen substrates (glcN). Different color lines show cumulative CO_2_ emissions (mean ± SD, n =4) derived from added glucose (light green) and soil organic matter (grey).

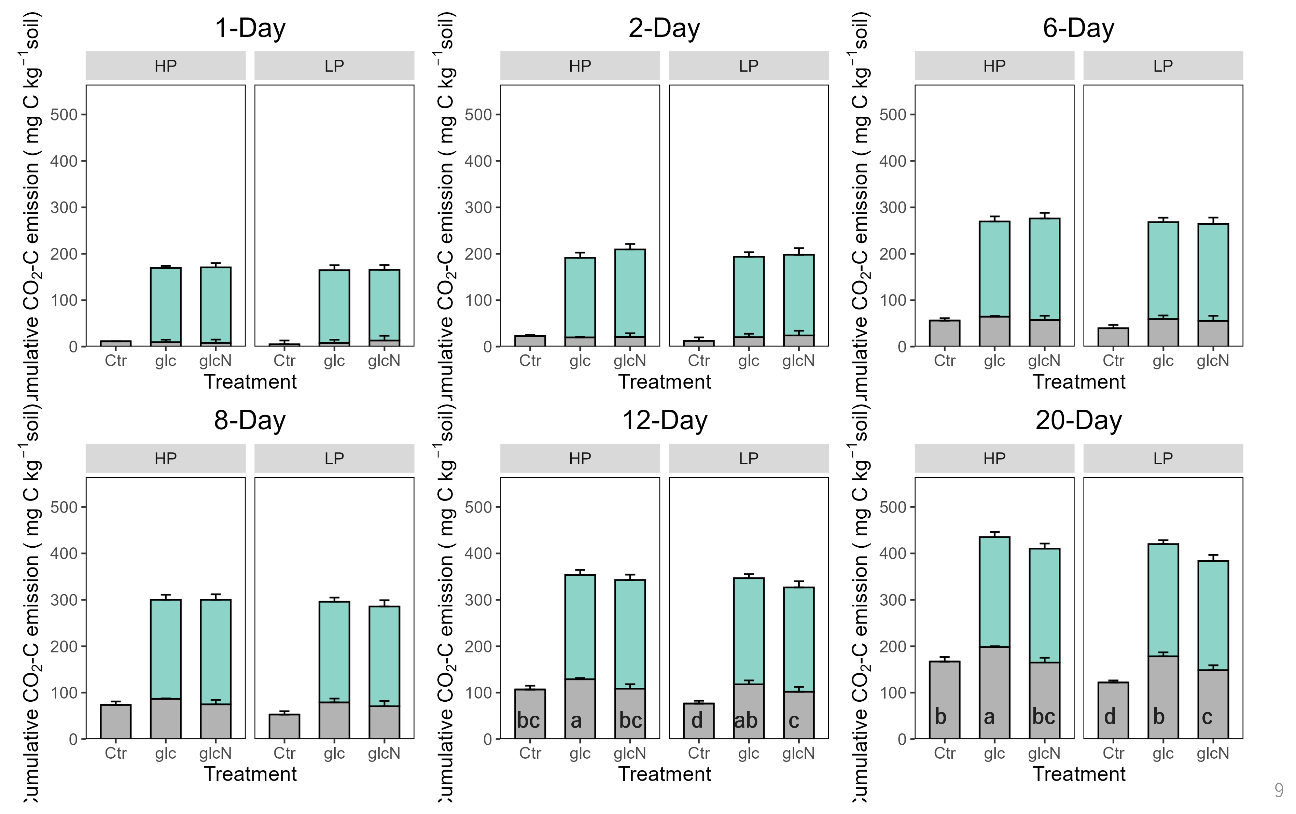

Fig S2. Cumulative CO_2_ emissions (mg C kg^–1^ soil) at 1, 2, 6, 8, 12, and 20 days after incubation. Treatments include soil without any amendment (Control), soil with ^13^C-enriched glucose (glc), and soil with ^13^C-enriched glucose and nitrogen substrates (glcN). Different color bars show cumulative CO_2_ emissions (mean ± SD, n =4, but n =3 in LP Ctr at 12 and 20 days due to an operational error during the experiment) derived from added glucose (light green) and soil organic matter (grey). Lowercase letters indicate significant differences (p <0.05) in SOM-derived CO_2_-C among treatments.
